# Burst Control

**DOI:** 10.1101/2021.02.13.431061

**Authors:** Eilam Goldenberg Leleo, Idan Segev

**Affiliations:** The Edmond and Lily Safra Center for Brain Sciences; Department of Neurobiology, the Hebrew University of Jerusalem, Israel

**Keywords:** Spike bursts, Dendritic computation, Single neuron computation, Coincidence detection, Calcium spike, NMDA spike, Dendritic inhibition, Synaptic plasticity, Nonlinear dendrites, Compartmental modelling

## Abstract

The output of neocortical layer 5 pyramidal cells (L5PCs) is expressed by a train of single spikes with intermittent bursts of multiple spikes at high frequencies. The bursts are the result of nonlinear dendritic properties, including Na^+^, Ca^2+^, and NMDA spikes, that interact with the ∼10,000 synapses impinging on the neuron’s dendrites. Output spike bursts are thought to implement key dendritic computations, such as coincidence detection of bottom-up inputs (arriving mostly at the basal tree) and top-down inputs (arriving mostly at the apical tree). In this study we used a detailed nonlinear model of L5PC receiving excitatory and inhibitory synaptic inputs to explore the conditions for generating bursts and for modulating their properties. We established the excitatory input conditions on the basal versus the apical tree that favor burst and show that there are two distinct types of bursts. Bursts consisting of 3 or more spikes firing at < 200 Hz, which are generated by stronger excitatory input to the basal versus the apical tree, and bursts of ∼2-spikes at ∼250 Hz, generated by prominent apical tuft excitation. Localized and well-timed dendritic inhibition on the apical tree differentially modulates Na^+^, Ca^2+^, and NMDA spikes and, consequently, finely controls the burst output. Finally, we explored the implications of different burst classes and respective dendritic inhibition for regulating synaptic plasticity.

## Introduction

Layer 5 pyramidal cells (L5PCs) are considered to be pivotal building blocks of the mammalian neocortex (DeFelipe and Fariñas 1992). In rodents, these cells receive ∼10,000 synaptic inputs over their dendritic surface, about 20% of them originate from the local microcircuit and the rest from external structures. The thalamus accounts for ∼10% of the synapses and represents the bottom-up input, while other cortical regions send connections reflecting top-down feedback and inter-module crosstalk (Rockland and Pandya 1979; Felleman and Van Essen 1991; Llinás 2002). The outputs of thin-tufted L5a PCs project laterally to nearby pyramidal cells and to other cortical areas. Thick-tufted L5b PCs project uniquely to subcortical regions (e.g., the thalamus; Z. Wang and McCormick 1993; Takahashi et al. 2020). Their key importance in processing information was already realized more than 100 years ago by Ramon y Cajal (1894), who termed cortical pyramidal cells “psychic” neurons.

Pyramidal cells in the neocortex *in vivo* tend to fire spikes in a random, Poisson manner such that the timing of each action potential is independent of its predecessors (Smith and Smith 1965). However, occasionally these cells also fire a brief burst of a few spikes at high frequency (Chagnac-Amitai, Luhmann, and Prince 1990). This occurs more than expected by chance (see **Methods**; Connors and Gutnick 1990). Specifically, deep layer 5-6 pyramidal cells tend to burst (Bastian and Nguyenkim 2001; van Elburg and van Ooyen 2010), and thick-tufted layer-5b pyramidal cells show both tonic and burst firing intermingled (Schwindt, O’Brien, and Crill 1997).

It was shown that spike bursts convey different information about stimuli compared to isolated spikes, or else serve some other specific function. In the primary visual cortex (V1) during drifting-gratings stimuli, spike bursts in putative pyramidal neurons were tuned to the spatial frequency and orientation of the grating, while isolated spikes were tuned to their contrast (Livingstone, M. S., Freeman, D. C., & Hubel 1996; Cattaneo, Maffei, and Morrone 1981). In the electric organ of weakly electric fish, single spikes encode self-position whereas bursts better represent communication with other individuals (Gabbiani et al. 1996). However, researchers continue to argue whether bursts generally encode different features than single-APs (Oswald 2004) or only sharpen the tuning (for review see Krahe and Gabbiani 2004).

Diverging lines of research probe burst involvement in additional functions other than feature-specific encoding. The BAC-firing coincidence detection mechanism (M. E. Larkum et al. 2009; M. Larkum 2013) employs burst firing to associate the activity of several presynaptic neurons impinging on different parts of the dendrite. Bursting also gives rise to a substantial increase in vesicle release probability at the synapse (Lisman 1997) and promotes switching between various states during sleep (Li, Poo, and Dan 2009). Bursts could allow for data multiplexing (Naud and Sprekeler 2018; Payeur, A., Guerguiev, J., Zenke, F., Richards, B., & Naud 2020), emphasize selective responses and propagate selective inputs (Balduzzi and Tononi 2013). Another debate still stands, whether more information is conveyed by the number of spikes in a burst (Eyherabide 2009; Csicsvari et al. 1998) or by their firing rate (Izhikevich et al. 2003). Other approaches probe burst relevance to plasticity, showing that pairing L5PC excitatory post-synaptic potentials (EPSPs) with bursts or with a single spike led to long-term depression (LTD) versus potentiation (LTP; Birtoli 2004).

At the biophysical mechanistic level, Larkum, Zhu and Sakmann (1999) were the first to demonstrate that bursting of cortical L5 pyramidal cells implements coincidence detection between perisomatic and tuft excitation (see also M. E. Larkum, Zhu, and Sakmann 2001; Schaefer et al. 2003; M. Larkum 2013). The basic *in vitro* procedure they used includes dual patch-clamp recordings targeting the soma and the main apical bifurcation (or ‘nexus’). In this paradigm, a backpropagating somatic action potential (bAP) lowers the threshold for a Ca^2+^ spike in the apical tree and, consequently, may lead to the generation of a somatic spike burst. They termed this phenomenon backpropagation activated Ca^2+^ spike firing (BAC firing). Many replicated and expanded their findings, including in some elaborate neuron models (Hay et al. 2011; Bahl et al. 2012). Without a bAP, dendritic input required for Ca^2+^ spike generation is much stronger and produces less somatic spikes (Matthew E. Larkum, Zhu, and Sakmann 1999). The lowest threshold for burst generation was found when tuft activation followed the somatic action potential by ∼5 ms.

There are several outstanding questions regarding the conditions for the initiation of bursts and the criteria for manipulating their characteristics. Details about the varied effects of dendritic inhibition on bursts are not yet clear. The necessity of the bAP for promoting Ca^2+^ spike firing and consequently bursting (Matthew E. Larkum, Zhu, and Sakmann 1999), is still questionable (Golding, Staff, and Spruston 2002). Furthermore, it is uncertain whether functional clusters of adjacent and temporally correlated inputs (whose existence was recently debated; Wilson et al. 2016; Chen et al. 2012) are sufficient for generating bursts?

The present work aims primarily to test various timing and location conditions of (basal and apical) excitatory synapses and of dendritic inhibition, for their effect on the initiation/control of somatic bursts. Towards this end, we employed the model built by Hay et al. (2011), that utilized an automated feature-based parameter search to faithfully replicate both dendritic Ca^2+^ and perisomatic Na^+^ electrogenesis, and the interaction between these two spiking regions (i.e., BAC-firing). Shai et al. (2015) employed this model to show how basal and tuft synapse numbers are transformed into high frequency bursts. This coincidence detection mechanism depends on voltage gated Ca^2+^ channels (VGCC), is approximated by a composite sigmoidal function, and is able to create orientation tuning. Our experiments introduce additional critical parameters that control this bursting mechanism, including dendritic inhibition, and suggest the involvement of bursts in Ca^2+^-dependent long-term synaptic plasticity.

## Results

### Temporal characteristics of excitatory dendritic inputs for burst generation

Spiking output of a cell depends critically on the number of activated synapses and their temporal correlation. To examine timing constraints on synaptic inputs for burst generation, we simulate a L5 PC model (Fig 1**a**; Hay et al. 2011; and see **Methods**), and look at the response as we vary activation times of a fixed suprathreshold number of synaptic activations (Fig 1**b**). The synapses, which incorporated AMPA- and NMDA-dependent conductances (see **Methods**) were uniformly distributed on the entire basal dendrite (blue region in Fig 1**a**), and on 25% of the apical tuft (750 μm continuous stretch, red region in Fig 1**a**; and see S2 Fig, and **Methods**). We first examined the minimal number of excitatory synapses over these dendritic subtrees that, when activated synchronously, elicit a burst of somatic Na^+^ spikes. We found a threshold for synchronous activation of all synapses with a peak AMPA and NMDA conductance of *g_max_* = 0.4 nS per synapse, at 50 ± 20 basal and 30 ± 10 apical tuft synapses.

**Fig 1.**
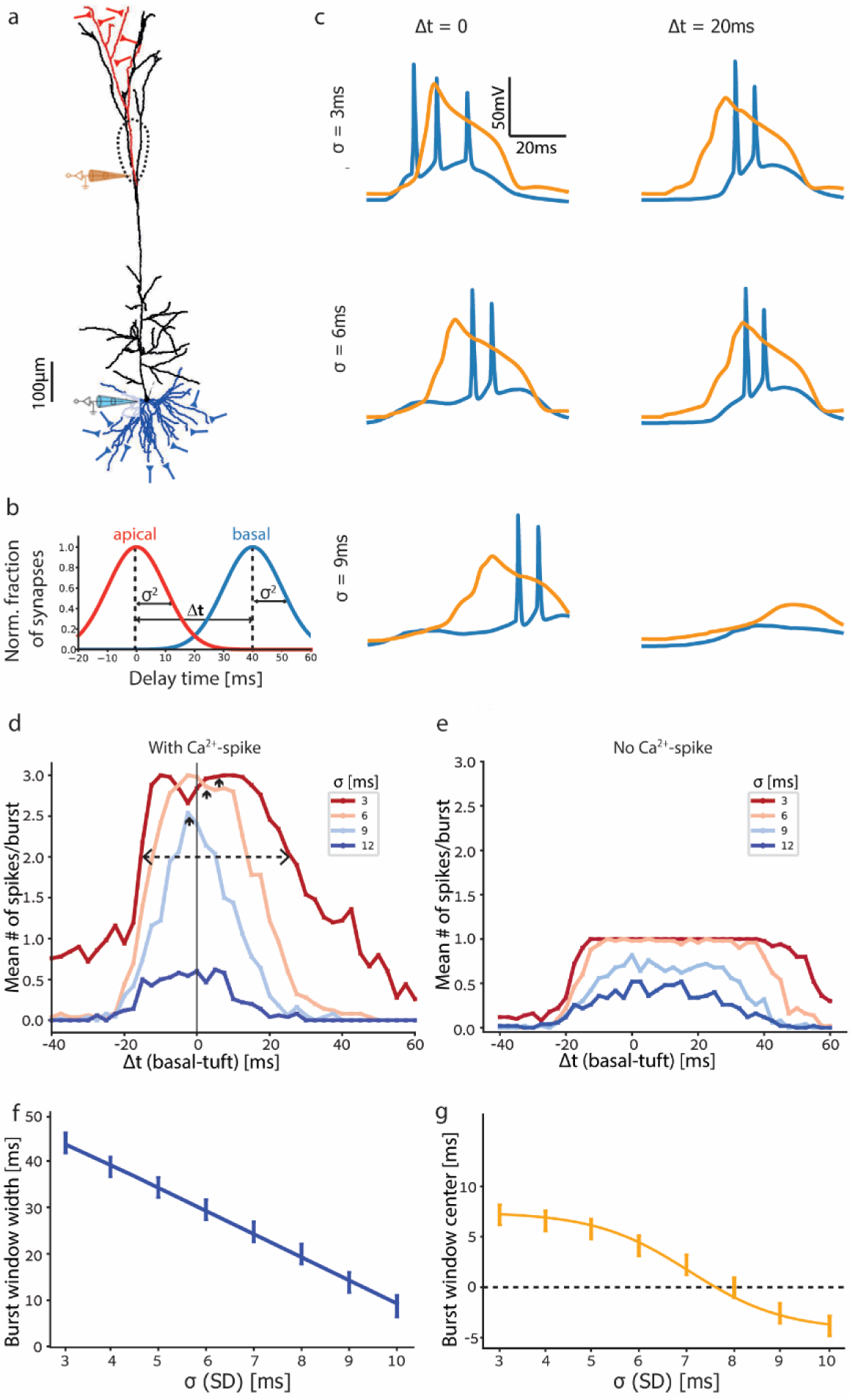
Temporal conditions for generating somatic spike bursts. **a**. Morphology of the modelled L5PC. Fifty excitatory synapses were distributed on the basal tree (blue synapses) and thirty on a subtree of the apical tuft (red synapses). Dashed line encircles the Ca^2+^ hotspot in the apical nexus. Blue and orange electrodes measure voltage traces shown in **c**. **b**. Two normal distributions of synaptic activation times for the basal and apical synapses are shown. The distributions have identical standard deviations, σ, and their means are shifted by Δt from each other. **c**. Voltage traces from the somatic and main apical bifurcation electrodes in **a** for a combination of σ and Δt values (Na^+^ spikes in blue and Ca^2+^ spike in orange). **d**. Mean number of somatic spikes per burst for a range of Δt values. Colored lines correspond to different σ values. The extent of Δt for generating a burst is reduced with increase in σ. Dashed line denotes curve width at 2 spikes per burst; arrows denote their centers. **e**. Same as **d** but without voltage-gated calcium channels, resulting in the absence of bursts. **f**, **g**. Width (**f**) and center (**g**) of the curves at two mean spikes per burst shown in **d**. Both width and mean reduce with increasing σ. Note in **g**, that the window’s center is positive (basal input following apical input) for smaller σ, and negative for larger σ.

Next, we explored the effect of the activation time of the synaptic input on burst generation. We randomly selected activation times for basal and apical synapses from two normal distributions of identical standard deviation σ, with a delay time of Δt between the mean activation times of these two sets of synapses (Fig 1**b** and S1**b** Fig). Typical voltage traces for different choices of σ and Δt are shown in Fig 1**c**. For Δt = 0 (left column in Fig 1**c**) raising σ values from 0 (instantaneous) to 3 ms did not change the output burst substantially (not shown). In both cases, the somatic burst comprised three Na^+^ spikes that were associated with a prominent Ca^2+^ spike (blue and orange traces respectively, in Fig 1**c** top). When increasing σ further to 6 ms and 9 ms (middle and bottom traces respectively, in Fig 1**c**), the burst consisted of only two spikes, whereas the Ca^2+^ spike remained essentially intact. Increasing the average delay Δt between the apical and basal synapses to 20 ms (right column in Fig 1**c**) with σ = 3 ms or 6 ms resulted in a burst of only two late spikes, and the dendritic Ca^2+^ spike remained intact. With further increase in σ to 9 ms, the Ca^2+^ spike was abolished and thus the somatic firing also terminated (lower trace in right column, Fig 1**c**).

The summary graph (Fig 1**d**) of the mean number of spikes per burst for a range of Δt and σ values, shows that more spikes/burst are fired for small values of σ and Δt. The range of Δt with > 2 spikes per trial on average (above the dashed line in Fig 1**d**) defines the conditions for burst generation. Specifically, for σ = 3 ms (red line in Fig 1**d**), the range of Δt for burst generation spans 40 ms from Δt = -15 ms (basal synapses preceding tuft synapses) to Δt = +25 ms (tuft synapses then basal synapses). Interestingly, this window is slightly biased towards Δt > 0 (tuft preceding basal activation), as evident from the center value marked by a small upwards arrow in Fig 1**d**. Increasing σ decreases the number of spikes per burst and the range of Δt giving rise to a burst (Fig 1**c** and 1**d**). For σ = 9 ms, the Δt window for burst generation narrows linearly to 13 ms (from Δt = -8 ms to +5 ms; Fig 1**f**). At this high σ, Δt values for burst generation are centered near -5 ms (basal input before tuft), which was found experimentally to have the minimum threshold of current injection for burst generation (Fig 1**d** and 1**g** as in Larkum, Zhu, and Sakmann 1999). At even higher σ = 12 ms, the mean number of spikes per burst dropped to 0.5 and bursting vanished (dark blue in Fig 1**d**). For control we ran this experiment again without voltage-gated Ca^2+^ channels at the Ca^2+^ hotspot. This condition results with 1-spike maximum and no bursting, as there is no Ca^2+^ spike to boost the tuft input (Fig 1**e**, compare to Fig 1**d**).

Stretching synaptic activation times using σ allows a proxy for testing lower input intensities and their requirements of tuft-to-basal delay for burst initiation. Increasing σ, monotonically decreased both the Δt range for burst generation and the Δt-window center (small upwards pointing arrows in Fig 1**d**). Increasing σ shifted the center from positive delays at low values (tuft input preceding the basal input) to small negative delays at larger values (Fig 1**g**). We conclude that broadening the timing of synaptic activation narrows the tuft-to-basal delay window for burst generation (Fig 1**d** and 1**f**) and shifts the preferred temporal order of activation from a tuft-then-basal to a basal-then-tuft sequence of activation (Fig 1**d** and 1**g**).

To verify the mechanisms for burst generation, we compared our model with the established backpropagation-activated Ca^2+^ spike firing paradigm (BAC; Matthew E. Larkum, Zhu, and Sakmann 1999) in S1 Fig. In this paradigm, a backpropagating somatic AP lowers the threshold for an apical Ca^2+^ spike and a burst. A somatic depolarizing current injection results in a single somatic AP. This AP backpropagates up to the apical tuft, where, if it meets a subthreshold current injection, they may together generate a dendritic Ca^2+^ spike and consequently a burst consisting of 3 APs at the soma. The somatic and dendritic (“nexus” main apical bifurcation) voltage traces in S1**c** Fig show responses to synaptic activations with variable σ at either or both basal and tuft dendritic trees. Our model reproduces BAC-firing using conductance-based synaptic activation alone on the dendritic trees (S1**c** Fig middle-row), instead of the original experimental and more artificial current injection (see Hay et al. 2011). For both the original clamp experiment and our simulation, stronger tuft input without perisomatic activation (achieved here by decreasing σ; S1**c** Fig bottom) generates a Ca^2+^spike (orange) and a burst of only two APs (blue; S1**c** Fig top-center).

To check for burst dependence on voltage-gated Ca^2+^ channels (VGCC), we set their conductance to zero at the Ca^2+^ hotspot and noticed that all spikes disappeared, except for a single somatic one (Fig 1**e** and S1**c** Fig right). Thus, we confirmed that low σ tuft input, combined with a somatic AP, creates a burst only via VGCC. Preserving VGCC and increasing σ to 9 ms caused tuft synapses to initiate a subthreshold voltage plateau, and prevented perisomatic synapses from generating an AP. Coincidental activation of the synapses on both trees may still elicit a full-blown Ca^2+^ spike and a somatic burst (S1**c** Fig bottom). Even larger σ > 10 ms with the fixed synaptic number will, at most, result in one somatic AP. We conclude that initiation of bursts occurs by either coincidental basal + tuft excitatory inputs, or by substantially strong excitatory input to the tuft alone.

### Spatial input conditions and temporal burst characteristics

We next explored the effects of the number of activated dendritic synapses that impinge on various parts of the dendritic tree, on burst generation (Fig 2). To do this we fixed the temporal parameters σ = 10 ms and Δt = 0, and separately manipulated the number of activated synapses on the basal tree and on a distal apical subtree (red subtree in Fig 1**a**).

**Fig 2.**
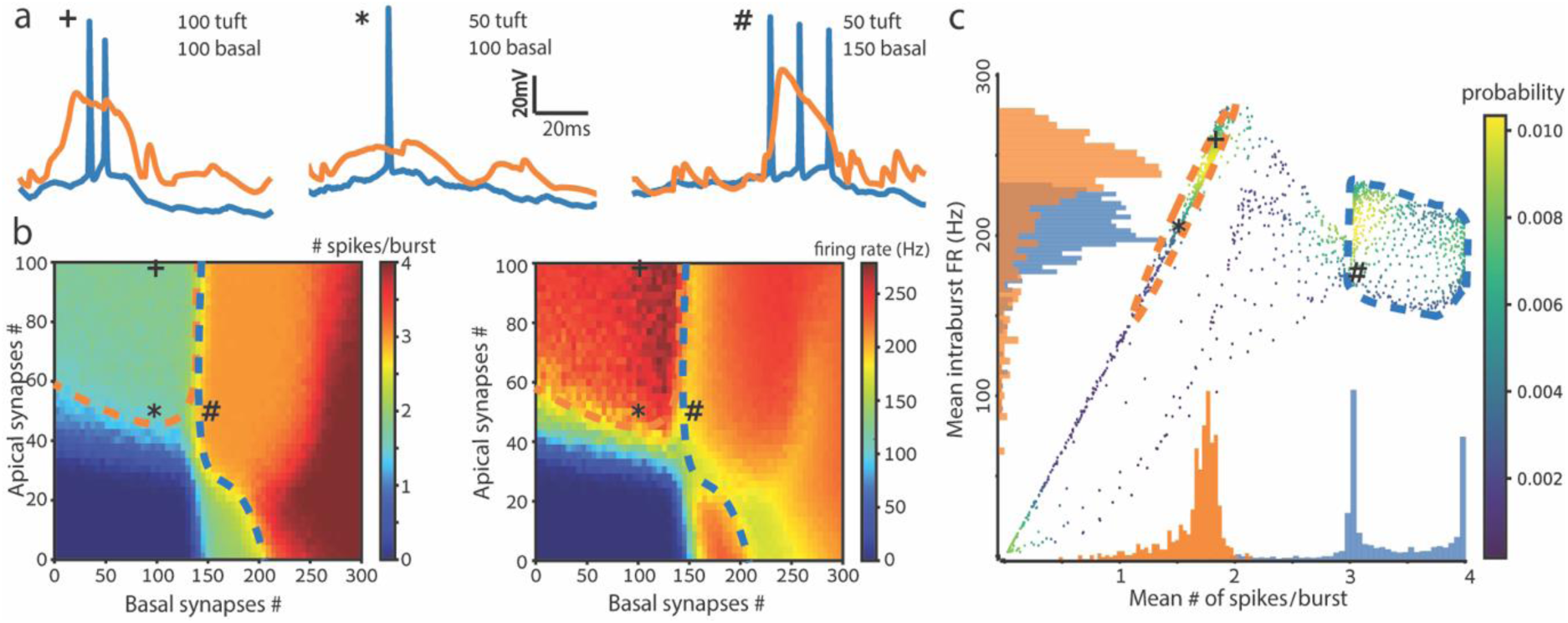
Two burst classes revealed based on the number of spikes per burst and intra-burst firing rate. **a**. Example somatic (blue) and dendritic (orange) voltage traces as in Fig 1 for the 3 cases of basal and tuft number of synapses: 100 and 100 (left), 100 and 50 (center), 150 and 50 (right). **b**. Heat maps of the mean # of spikes (left) and the intra-burst firing rate (right) for a range of numbers of basal and tuft synapses. The bursting threshold (> 2 spike/burst blue-to-green transition) is 40-60 tuft synapses or 140-150 basal synapses. Dashed lines denote smoothed class borders. Bursts with 3-4 spikes appear only for strong basal input (within the blue dashed lines). Apical tuft input above threshold results in shorter bursts of 2 spikes and higher rates (within the orange dashed lines). **c**. Scatter plot for each combination of synapses (x-y coordinates) in **b**, sorted by the mean # of spikes per burst (left frame) and the intra-burst firing rate (right frame). Each value in right-hand heat map is plotted with its corresponding value in the left-hand heat map. The color of each point in **c** represents the density of data points with the same values, equivalent to their probability. The orange and blue histograms along the x- and y-axes correspond to the data in the two clusters encircled in orange and blue, respectively. Temporal input parameters were fixed at Δt = 0, σ = 10 ms.

Fig 2**a** depicts three example voltage traces. When 100 basal and 100 tuft excitatory synapses were activated, a dendritic Ca^2+^ spike was initiated, accompanied by a burst of 2 somatic spikes (orange and blue respectively, Fig 2**a**, left). When the number of activated tuft synapses was reduced to 50, the burst disappeared and so did the dendritic Ca^2+^ spike (Fig 2**a**, center). Keeping the number of activated tuft synapses at 50 but increasing the activated basal synapses to 150 resulted in a burst of 3 spikes (with lower FR, see below) accompanied by a dendritic Ca^2+^ spike (Fig 2**a**, right).

Next, we measured the mean number of spikes per burst while independently varying the number of activated synapses impinging on both the basal dendritic tree and the apical tuft. Activating up to 50 apical and 140 basal synapses (with Δt = 0, σ = 10 ms) is mostly insufficient for generating a burst, and produces up to one spike (blue regions, Fig 2**b**; “*” indicates the example trace 2**a** center). 60 ± 10 apical synapses without basal synapses are sufficient for the generation of a burst of two spikes (top green region, Fig 2**b** left; “+” denotes the example trace, 2**a** left). A burst of two spikes is observed when 150 - 200 basal synapses (without apical synapses) are activated; increasing the number of basal synapses to 200 - 300 results in bursts of 3 or 4 spikes (orange and red regions, respectively, Fig 2**b** left; “#” denotes the example trace 2**a**, right). This finding is counterintuitive as tuft synapses are thought to be less potent in generating somatic spikes, due to the attenuation of their effect along the ∼1 mm distance of the tuft dendrites from the soma. However, tuft synapses compensate more for this attenuation by NMDA spike generation, allowing burst-firing by fewer synapses (see **Discussion**).

A supplementary analysis is shown in S3 Fig where the apical:basal ratio of the number of synapses is varied while the total number of activated synapses is fixed. For 50 or 100 total activated synapses, apical synapses alone promoted firing of one or two spikes, respectively, whereas the same numbers of basal synapses do not generate any spiking (red and grey lines in S3 Fig). 200 apical synapses alone generate bursts of 2 spikes, less spikes than with 200 basal synapses (2.5 spikes) or with 50 apical and 150 basal synapses (3 spikes/burst; blue line in S3 Fig).

Fig 2**b** right shows the intra-burst firing rate as a function of the number of basal and tuft synapses. An apical input of > 50 activated synapses, with a basal input of < 150 synapses, generates high-frequency bursts of ∼250 Hz or, equivalently, inter spike interval (ISI) ≈ 4 ms (dark red area within orange dashed line in Fig 2**b** right), whereas higher basal input intensity of > 150 synapses results in bursts with lower rates of ∼200 Hz (orange region, right of the blue dashed line, Fig 2**b** right). Complementing Fig 2**b** left, our findings reveal a separation of bursts into discreet classes. One class of bursts contains a small number of spikes at high rates (2-3 spikes at ∼250 Hz); this class is generated by the activation of many apical tuft synapses and mediated by an NMDA spike. The other class of bursts contains a larger number of spikes fired at lower rates (3-4 spikes at ∼200 Hz) and is promoted by detection of coincidence of basal input with weak apical tuft activation.

Fig 2**c** is a scatter plot that combines the results of both plots in Fig 2**b**, and shows the mean number of spikes per burst (x-axis) versus the intra-burst firing rate (FR; y-axis). Namely, each point in Fig 2**c** represents the number of spikes per burst from Fig 2**b** left frame and its respective FR from Fig 2**b** right frame, for the same basal and apical synapse numbers. Three clusters emerged at regions of yellow and green points in Fig 2**c**, which are colored according to the probability of both spikes/burst and FR values. One cluster formed on the upper linear line in the scatterplot (orange dashed line, Fig 2**c**). It is characterized by a high FR of ∼250 Hz and a mean of two spikes per burst. We believe this burst class is driven by an NMDA spike (see **Discussion**). The second cluster appears as a rectangle to the right and is characterized by bursts of three or four spikes at a lower FR of ∼200 Hz (blue dashed line, Fig 2**c**). It is associated with coincidence detection of input to the basal dendrites and to the apical tuft, resembling BAC-firing as suggested by Larkum et al. (1999). The third cluster is located near the origin and reflects the case where no spikes were initiated, corresponding to the subthreshold blue areas in the two frames of Fig 2**b**.

A further quantification is attained by superimposed histograms of both burst classes (Fig 2**c**): in orange the high FR and two spikes class, and in blue the lower FR and 3-4 spikes class (see **Methods**). These reveal clear separation in spike number and partial discrimination in intra-burst FR (histograms on x and y axes, respectively, Fig 2**c**). The scatterplot transforms the input-aligned picture of the heatmaps into a map of output burst characteristics showing each mean spikes-per-burst value with its corresponding FRs. Accompanied by the associated histograms, the scatterplot further elucidates the clear separation between the two burst classes.

### Editing bursts with inhibition

A local modulation of pyramidal cell activity is provided by a spectrum of inhibitory interneuron types that are distributed over its dendritic tree (DeFelipe and Fariñas 1992). We tested the sensitivity of burst generation to different locations of inhibitory synapses along the apical tree (Fig 3). Fig 3 shows a burst of 3-spikes, and the various edits that targeted inhibition may perform on it for different inhibition locations. We first selected a single apical branch (Fig 3**a**) to receive excitatory synapses. Inhibition was introduced by activating 20 GABA_A_ synapses at one of the numbered dendritic locations at each experimental condition (#1 - #9, Fig 3**a**). Varying the location of the activated inhibitory synapses produced the respective voltage traces in Fig 3**b**. The control case without inhibition is shown in Fig 3**c**.

**Fig 3.**
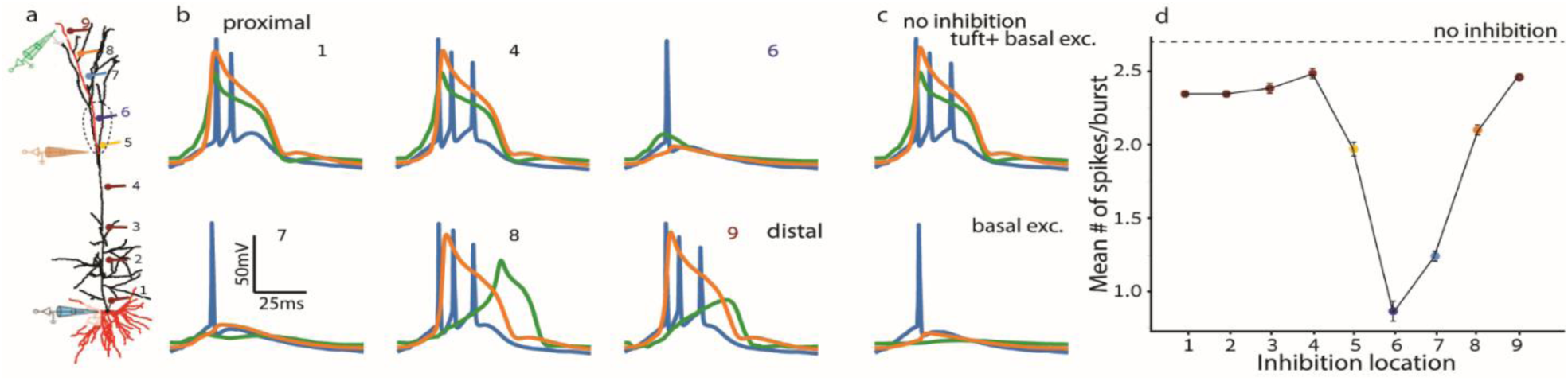
Inhibitory synapses edit the output burst differently depending on their dendritic location. The case of excitatory and inhibitory synapses restricted to a part of the apical tree. **a**. Model layer- 5b thick-tufted pyramidal neuron as in Fig 1a. Electrodes indicate locations of voltage recording. Red branches in both basal and apical trees receive distributed excitatory synapses. **b**. Voltage traces from a somatic (blue) and two dendritic electrodes (orange and green) shown in **a**, during the activation of inhibition in a single locus (1-9) in the apical trunk (1,4) and apical tuft (6-9), indexed as that of the synaptic location in **a**. Proximal dendritic inhibition at location (1) suppresses the first Na^+^ spike (blue trace; compare to **c**, top trace) whereas inhibition at locations (6,7) abolishes the Ca^2+^ spike (orange) and NMDA spikes (green) and, consequently, the latter two spikes in the burst. Distal inhibition (8-9) only attenuates the terminal-branch NMDA spikes (green trace) but does not significantly affect the Ca^2+^ spike (orange trace) nor the somatic burst. **c**. Voltage traces without inhibition for basal and tuft excitation (top) and for basal-only excitation (bottom) for comparison. **d**. Mean # of spikes per burst as a function of location of inhibitory synapses (as in **a**). In all cases 70 basal and 40 tuft excitatory synapses were activated simultaneously (Δt = 0) with σ = 10 ms (see **Methods**). For simulating inhibitory input, 20 inhibitory synapses were uniformly distributed up to 100 μm from each location (1-9) as marked in **a**; the peak conductance per inhibitory synapse was 1 nS (see **Methods**).

When compared to the control case without inhibition (Fig 3**c**), we found that perisomatic apical trunk inhibition (at location #1) suppresses the first somatic spike (blue trace in Fig 3**b**1) but does not affect the dendritic Ca^2+^ spike (orange trace) nor the NMDA spike in the apical tuft (green trace). Distal trunk inhibition does not affect the Ca^2+^ spike, nor any somatic spikes (compare Fig 3**b** location #4 to Fig 3**c** top – no inhibition). Inhibition centered at the Ca^2+^ hotspot (#6) or adjacent to it, at about 100 μm distal (“off path”; #7) suppressed the Ca^2+^ spike and, consequently, the two latter somatic Na^+^ spikes, thus abolishing the burst. The impact of “off path” (rather than “on path”) inhibition on dendritic excitability was demonstrated and discussed by Gidon and Segev (2012). Finally, inhibition acting on the distal tuft hardly modulates Ca^2+^ or somatic Na^+^ spiking (compare locations #8 and #9 to Fig 3**c** top), but only alters local NMDA spikes (green traces).

The effect of the dendritic location of inhibitory synapses on burst generation is summarized in Fig 3**d**, showing the mean number of somatic spikes per burst for each inhibition locus (numbered as in Fig 3**a** and 3**b**). Clearly, the most effective inhibition on burst generation is located 200 – 400 μm distal to the apical nexus (points #6 and #7, respectively). We varied the temporal order and separation (Δt_inh_) of excitation and inhibition and found no differential effect between dendritic locations, except for the effective time window width for inhibition. The inhibition with the largest effect (in location 6) disrupts spiking when activated even ±15 ms before or after excitation. Inhibition with a smaller effect (at locations #5, #8) contributes only when activated synchronously with excitation (not shown).

Next, we tested the case where the excitatory synapses were distributed over the entire apical tuft. Inhibitory synapses were distributed at all branches with a fixed distance from the soma (Fig 4**a**). The traces in Fig 4**b** show the influence of inhibition on somatic (blue) and dendritic tuft (orange red and green) voltage during burst generation. Each row relates to a single distance of inhibitory synapses: 100 μm from soma in bottom row, 900 μm in center, and 1100 μm in top. Each column depicts inhibition at a different timing condition: left Δt_inh_ = 0, center Δt_inh_ = -10 ms (inhibition before excitation), and right Δt_inh_ = +10 ms (inhibition after excitation).

**Fig 4.**
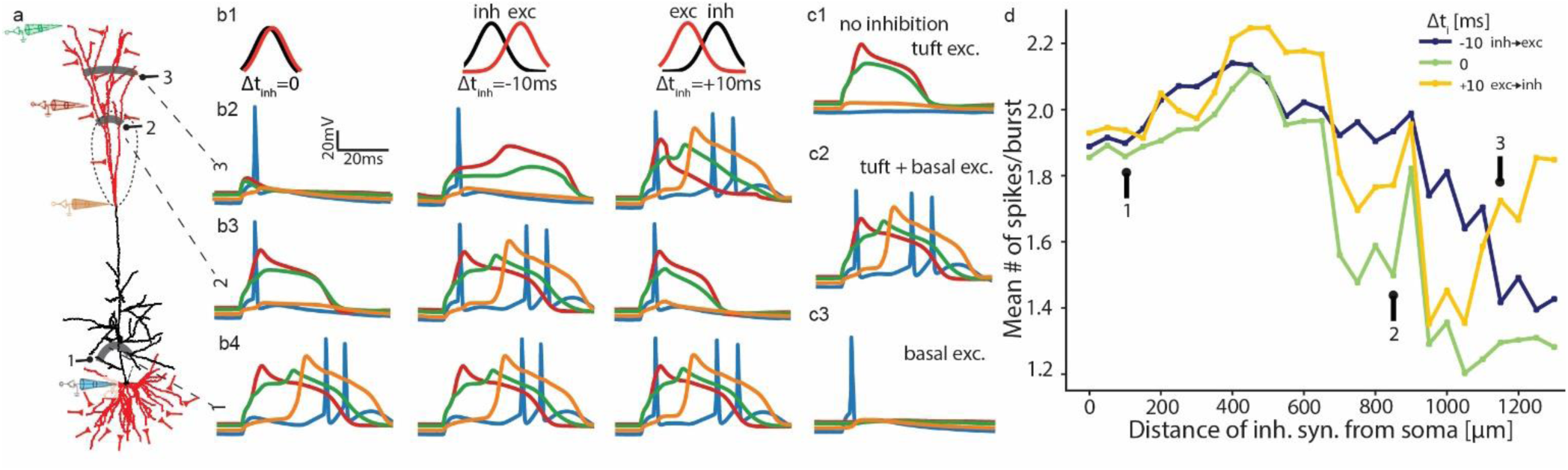
Impact of dendritic inhibition on burst generation. The case of whole apical tuft activation with dendritic inhibition distributed at strips of iso-distance from soma. **a.** Schematics showing the distribution of synapses. Inhibitory synapses at all locations of a fixed distance from the soma (a single numbered grey shaded strip). Synapses #1 at 100 μm from the soma on the oblique branches; synapses #2 at 900 μm from the soma on the intermediate apical tuft and synapses #3 at 1100 μm on the distal apical tuft. Electrodes correspond to colored voltage traces in **b**. **b1.** Normalized distribution of synaptic activation times separated for excitatory and inhibitory inputs, with different delays Δt_inh_ for the three columns shown. **b2-b4.** Voltage traces for inhibition at distances 1, 2 and 3, respectively. **c1-c3.** Control cases. **c1.** Forty excitatory synapses activated at the tuft triggers NMDA spikes but no dendritic Ca^+2^ spike nor somatic spikes. **c2.** Excitatory input to both basal (70) and tuft (40) dendrites generates an NMDA- (red/green) and a Ca^2+^ spike (orange) and, in turn, a somatic burst of Na^+^ spikes (blue). **c3.** Seventy excitatory synapses activated at basal dendrites generate a Na^+^ spike. **d.** Mean # of spikes in a burst as a function of the distance of inhibition from the soma. Lines correspond to different Δt_inh_ values as in **b**1. In all cases, the excitatory synapses are as in Fig 3 (70 basal and 40 tuft), but distributed on the entire tuft, activated simultaneously (Δt = 0) with σ = 10 ms (see **Methods**). In each inhibitory stripe shown in **a**, 20 inhibitory synapses were activated in a 200 μm uniform distribution around all branches at each distance from the soma (1-3).

Fig 4 shows that for 70 basal and 40 diffuse apical excitatory synapses, diverging inhibition achieves similar efficiency to the single-branch case (compare Fig 3**b** to Fig **4b** left Δt_inh_ = 0, and compare Fig 3**d** to Fig **4d** green): Perisomatic inhibition (location 1 in Fig 4**a**) suppresses the first spike of the burst (blue traces in Fig 4**b** bottom); above the Ca^2+^ hotspot (location 2) inhibition eliminates the Ca^2+^ spike (orange in Fig 4**b** center row; compare to Fig 4**c**2 – no inhibition), and at distal terminals (location 3) it attenuates close-by NMDA spikes whose absence disrupts the Ca^2+^ spike and burst firing (red and green traces in Fig 4**b** top; compare to Fig 4**c**1 – tuft excitation alone).

Surprisingly, this experiment reveals a novel inhibitory time dependence (Fig 4**b** columns and 4**d**), by which all the aforementioned inhibitory consequences appear at Δt_inh_ = 0 (Fig 4**b** left column), and at either Δt_inh_ = -10 ms (center column; inhibition before excitation), effective perisomatic (location #1) and distal inhibition (#3), or at Δt_inh_ = +10 ms (right column; inhibition after excitation), effective inhibition around the Ca^2+^ hotspot (#2). Only this last delayed inhibition effect resembles modelling and experimental results (Doron et al. 2017; Du et al. 2017). A comparison between inhibition effectiveness at various locations and under these three timing conditions (Fig 4**d**) exhibits preference for concurrent inhibition (Δt_inh_ = 0; green), then inhibition following excitation (Δt_inh_ = +10 ms; yellow) in intermediate tuft and preceding excitation (blue) in distal terminals.

The blue traces in Fig 4**b** show somatic spiking (intra-burst spike-count lower for both proximal trunk and tuft inhibition, see Fig 4**d**), and some additional depolarization from perisomatic (basal) current sources. Orange traces signify the Ca^2+^ spike at the hotspot (above the nexus) occurring for all tuft activations except nexus-proximal tuft inhibition (Fig 4**b** central row; decoupling NMDA spikes from the hotspot), either after voltage accumulation during extensive NMDA spikes (right) or concurrently with a somatic bAP (left). Red traces exhibit mid-branch NMDA spikes, and green their distal parallels, both eliminated by distal inhibition (Fig 4**b** top row). Each spike form and locus are manipulated by inhibition at a location ineffective for the other spike types, and only in part of the delay conditions.

Proceeding with whole-tuft excitation configuration, but slightly changing the balance of excitatory synapses between dendrites to 80 basal and 30 apical synapses (from 70 and 40 in Fig 4) creates a relatively “tuft-independent” burst generating scheme, where even strong (20 nS) tuft inhibition distributed on all branches of any one fixed distance from the soma (and nexus) does not prevent initiation of a Ca^2+^ spike and a burst. However, bursting is here affected by inhibition at the mid-upper apical trunk, decoupling the Ca^2+^ hotspot from its igniting bAP which did not happen in Fig 3 or Fig 4 (not shown).

We determined optimal inhibition spatial extent by adjusting the uniform synaptic distribution to a variable portion of the apical tuft, the same as previously described for excitation, and presented together (S2 Fig). The optimal spatial dispersion for 20 inhibitory synapses with 1 nS peak conductance each (see **Methods**), which maximally decreases the number of spikes per burst, was 2.5-5% of the total apical dendritic length (5-10% of tuft length), or 185-370 μm. As illustrated in S2 Fig, this dispersion is at half or less the optimal extent of excitatory synapses (measured inversely, by finding maximal spikes per burst). We therefore fixed the dendritic length containing inhibition at 200 μm, i.e., a mean synaptic density of one inhibitory synapse in every ∼10 μm, in line with experimental data (Chen et al. 2012), thus determining the specific locations most susceptible to inhibition of bursting.

### Impact of inhibiting bursts on the plasticity of excitatory synapses

Recent studies associate inhibition of bursts and of dendritic Ca^2+^ spikes with restriction of plasticity to specific connections for efficient learning (Owen, Berke, and Kreitzer 2018; Grienberger, Chen, and Konnerth 2014; Pérez-Garci, Larkum, and Nevian 2013). To study how inhibition of bursts is associated with synaptic plasticity in the apical tuft, we utilized an established Ca^2+^-dependent plasticity model (Shouval, Bear, and Cooper 2002; Bar-Ilan, Gidon, and Segev 2013; Eqs. (1) and (2) in **Methods**, and Fig 5a1, 2). Low Ca^2+^ concentration, [Ca^2+^]_i_, values result in “protected” (unchanged) efficacy of the excitatory synapses (Fig 5**a**, PROT.), higher [Ca^2+^]_i_ produce long-term depression (LTD, Fig 5**a**1, blue), and even higher values elicit potentiation (LTP, Fig 5**a**1, red; Shouval, Bear, and Cooper 2002). A Ca^2+^- dependent learning rate, η, (Fig 5**a**2) multiplies changes in efficacy (Eq. (2) in **Methods**) to fit experimental findings. Using our model, we tested how L5PC excitatory synapses act under this plasticity rule, and explicitly how inhibition of dendritic Na^+^, Ca^2+^ and NMDA spikes (determined by location and Δt_inh_) controls the manifestation of this plasticity rule.

**Fig 5.**
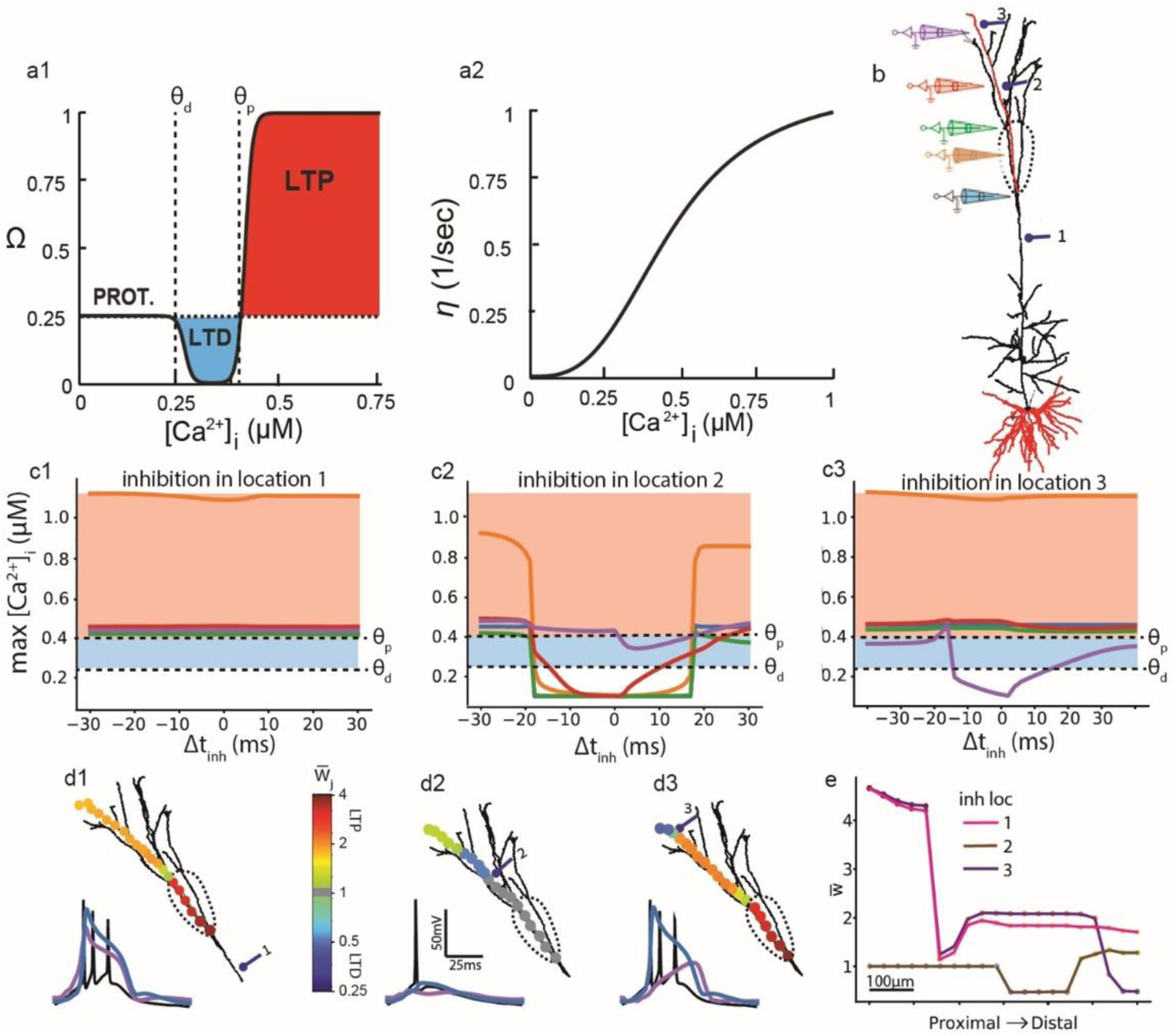
Impact of dendritic inhibition on somatic bursts and on plasticity of excitatory dendritic synapses. **a1**. Dependence of plasticity function, Ω, on intracellular Ca^2+^ concentration, [Ca^2+^]_i_. When [Ca^2+^]_i_ < θ_d_, the synaptic weights remain fixed (PROT. = protected); for θ_d_ < [Ca^2+^]_i_ ഼ θ_p_, the synapse undergoes long-term depression, LTD (blue) and for [Ca^2+^]_i_ > θ_p_ it undergoes long-term potentiation, LTP (red), see Eq. (1) in **Methods**). **a2**. Learning rate, η, as a function of [Ca^2+^]_i_ (adapted from Bar-Ilan, Gidon, and Segev 2013; see Eq. (2) in **Methods**). **b**. Model neuron and synaptic simulation parameters as in Fig 3a. 70 basal and 40 tuft excitatory synapses were activated (the “input”) on the red branches simultaneously (Δt = 0) with σ = 10 ms (see **Methods**). Electrodes and their respective color represent measurement locations of [Ca^2+^]_i_. Numbered synapses correspond to the mean locations of 20 inhibitory synapses with peak conductance 1 nS per synapse. Dashed line denotes Ca^2+^ hotspot. **c1-c3**. Maximal [Ca^2+^]_i_ along the apical dendrite at five locations (denoted by the colored electrodes in **b**) following the input as a function of timing, Δt_inh_, between excitation and inhibition, for each inhibitory location (#1-#3 in **b**). Dashed lines mark threshold values, θ_p_ and θ_d_, as in **a**1. Blue shading - LTD; red - LTP; white - protected. **d1-d3**. Up: Twenty representative excitatory synapses are superimposed on the dendritic tree, color coded by their respective synaptic weights at each inhibitory location, #1-#3, following the plasticity protocol with 10 input repetitions for the case of Δt_inh_ = 0. Lower traces: Somatic (black); nexus Ca^2+^ spike (blue trace at respective blue electrode in **b**) and distal NMDA spike (purple at respective purple electrode in **b**). for inhibition location #1. **d2**. As in **d**1, with inhibition at location #2 - burst and Ca^+2^ spike suppressed. **d3**. As in **d**1, with inhibition at location #3 - distal NMDA spike suppressed, burst is unaffected. **e**. Change in synaptic weights following the plasticity protocol as a function of synapse location along the tuft branch, for each of the three inhibitory locations.

Fig 5**b** shows the modelled tree and the locations of the inhibitory synapses and measuring electrodes. Synaptic configuration is identical to that used in Fig 3. 70 basal and 40 tuft excitatory synapses were activated with (excitatory) Δt = 0 and σ = 10 ms. Fig 5**c**1-3 plots the maximal [Ca^2+^]_i_ at each electrode position (lines correspond to colored electrodes in Fig 5**b**) for a range of Δt_inh_ values (x-axis) with trunk (1), intermediate tuft (2) or distal (3) inhibition. Note that this maximum value of [Ca^2+^]_i_ does not fully represents whether an excitatory synapse at the location will be potentiated or depressed, as the time that [Ca^2+^]_i_ lasts at any given concentration will eventually determine the sign of plasticity.

Fig 5**c**1 depicts the case in which the inhibition is proximal to the Ca^2+^ hotspot (at location #1) on the apical trunk. The maximum [Ca^2+^]_i_ at the hotspot is high and nearly invariant to inhibition timing, (orange line in Fig 5**c**1), resulting with LTP for all Δt_inh_ values for synapses at the hotspot. The maximum [Ca^2+^]_i_ at the more distal locations is lower but above the potentiation threshold θ_p_, so weaker LTP is observed there (Fig 5**c**1). Fig 5**d**1 depicts the same case, for Δt_inh_ = 0, by superimposing the excitatory synapses on the dendritic tree, color coded by their respective synaptic weights. Ten input repetitions were introduced in succession for a magnified effect on synaptic weights. For this proximal inhibition at location #1 the somatic burst is unaffected and so is the Ca^+2^ spike at the hotspot (blue trace) and the distal NMDA spike (purple trace).

Fig 5**c**2 and 5**d**2 depicts the case in which the inhibition is 100 μm distal to the Ca^2+^ hotspot (at location #2). Now, the synapses proximal to inhibition including the hotspot are “protected”, and distal to it undergo LTD; only for the large positive or negative Δt_inh_ values do synapses undergo LTP (Fig 5**c**2). In this case for Δt_inh_ = 0, the somatic burst is suppressed and so does the Ca^2+^ spike at the hotspot as well as the distal NMDA spike (Fig 5**d**2). Finally, Fig 5**c**3 and 5**d**3 depict the case in which the inhibition is at distal location #3. Now the proximal synapses undergo LTP whereas synapses distal to the inhibition undergo LTD. In this case, the somatic burst and the Ca^2+^ are unaffected whereas the distal NMDA spike is suppressed. A summary graph of excitatory efficacy changes at all dendritic locations for each inhibition location and Δt_inh_ = 0 is presented in Fig 5**e**. Note that the synaptic efficacy following the plasticity protocol are highly sensitive to the choice of threshold values for plasticity. In Fig 5 we fixed the thresholds at values conforming with previous modelling studies; in the supplementary analysis we provide additional results when varying θ_p,d_ (S4 Fig).

We briefly summarize the key results of this section by noting that (1) inhibition located at the hotspot or immediately distal to it suppressed Ca^2+^ spiking and branch-wide LTP; (2) inhibition of local NMDA spikes controls branch-specific plasticity by transforming LTP to local LTD or protection, and (3) following the association of bursts, Ca^2+^ spikes and inhibition with plasticity (see **Discussion**), we find a correlation between inhibition of bursts and limiting plasticity.

Finally, we tested for diverse outcomes of plasticity by the different burst classes and their corresponding inhibition, by utilizing the same Ca^2+^-dependent plasticity rule with whole-tuft excitation as in Fig 4**a**. Full tuft excitation was mainly associated with low [Ca^2+^]_i_, meaning minor plasticity effects in the burst-suppressed dendritic tuft (S5 Fig). With these final results, our plasticity findings ultimately connect temporal (Fig 1 and S1 Fig) and spatial (S2 Fig) conditions for bursting with class separation (Fig 2), now evidently providing a functional basis (NMDA-burst dependent plasticity), through differential inhibition (Fig 3 and Fig 4) and an implementation for restricting and tuning Ca^2+^- dependent plasticity (Fig 5 and S5 Fig).

## Discussion

Several studies have demonstrated the importance of spike bursts in pyramidal neurons in neural coding schemes (Krahe and Gabbiani 2004), plasticity (Royer et al. 2012; Goldberg, Lacefield, and Yuste 2004), and information flow (M. Larkum 2013; Lisman 1997). However, the input conditions for burst generation in the single cell are not fully understood, and neither are the effects of dendritic inhibition on shaping burst properties and on synaptic plasticity in dendrites. This work was aimed at filling these gaps, using a biologically faithful model of a layer 5 thick-tufted pyramidal cell.

### Excitatory conditions for burst generation – two types of bursts

We showed that somatic bursts arise either from a bidirectional stream of excitatory inputs combining basal and tuft synapses (Fig 1), or by generation of a local dendritic voltage plateau due to clustered excitatory input to the apical tuft alone (Fig 2 and S1 Fig). Bursts that result from a coincidence of basal and apical synaptic activation consist of 3-4 spikes fired at a frequency of ∼200 Hz (Fig 2), whereas the activation of > 30 excitatory synapses on a length of 350 μm – 750 μm of the dendritic tuft (S2 Fig) generates NMDA spikes at several apical loci (Alon Polsky, Mel, and Schiller 2004) initiating a Ca^2+^ spike at the main tuft and a high-frequency burst of ∼2 spikes. The number of tuft synapses required for burst generation is less than half that of basal synapses (Fig 2 and S3 Fig). This finding is counterintuitive, as tuft synapses are thought to be less potent in generating somatic spikes, due to steep voltage attenuation expected in a length of ∼1 mm along the apical dendrite towards the soma. Several factors compensate for the attenuation and make tuft synapses more likely to than basal synapses initiate bursts: clustered distribution (S2 Fig), high excitability of thin tuft terminals (Alon Polsky, Mel, and Schiller 2004), and proximity of tuft synapses to the Ca^2+^ hotspot which promotes generation of Ca^2+^ spikes. Further experiments should be conducted to explore whether these two burst types exist in cortical L5 pyramidal neurons, and to reveal the implications of the different types for neuronal computation and for plasticity-related processes (see below).

In agreement with Larkum et al. (1999), we found similar conditions on the timing of synaptic activation for burst generation when basal-then-apical (Δt < 0) synapses are activated (Fig 1). However, by varying the standard deviation, σ, of the apical and basal synapses activation times (Fig 1 and S1 Fig) we found that, for small σ (< 10 ms) burst generation is also enabled in the reverse tuft-to-basal order of activation (0 ms < Δt < 30 ms; Fig 1). This resulted from the long timescales and boosting effect of NMDA potentials that benefited from the bAP’s arrival after initial voltage buildup at the apical dendrites. This prediction still awaits experimental validation.

### Inhibitory control of bursts

Next, we examined how the location and timing of dendritic inhibition ‘edits’ the somatic bursts. To the best of our knowledge, this is the first direct theoretical study of this question. For excitation on a single tuft branch, perisomatic apical trunk inhibition (e.g., via basket cells) abolished the first somatic spike but left the dendritic spikes unchanged and allowed for burst initiation (Fig 3**b**1). Inhibition in or immediately distal to the Ca^2+^ hotspot (“off path” condition; see Gidon and Segev 2012) disrupted the Ca^2+^ spike and suppressed the somatic burst (Fig 3**b**6-7 and 3**d**), whereas more distal inhibition attenuated local NMDA spikes without affecting the burst (Fig 3**b**8-9).

A burst induced by whole-tuft excitation was suppressed by inhibition distributed at all branches of a fixed distance distal and adjacent to the hotspot, for Δt_inh_ ≥ 0 (inhibition following excitation; Fig 4). Δt_inh_ ≤ 0 is more effective in disrupting burst generation when inhibition is activated in distal locations, by suppressing NMDA spikes there. Contrary to our results, theoretical and experimental studies found the most effective timing of inhibition for suppression of NMDA spikes to be after excitation (Doron et al. 2017; Du et al. 2017). This is because they use basal activation and set the criterion for efficiency of inhibition as the reduction in the voltage integral, rather than the strong instantaneous onset of NMDA spikes that is crucial for burst formation. Overall, we showed that by controlling local dendritic excitability, in particular the NMDA and/or Ca^2+^ spikes, local dendritic inhibition can finely edit the output bursts.

### Effect of inhibition on dendritic plasticity – relationship to burst control

Notably, intracellular Ca^2+^ concentration is implicated in long-term synaptic plasticity (Rose and Konnerth 2001; Zucker 1999). Utilizing the calcium-dependent synaptic plasticity model (Shouval, Bear, and Cooper 2002), our study showed that inhibition, during somatic bursts and correlated Ca^2+^ spikes, enables diverse plasticity modification maps. Due to well-located and timed dendritic inhibition, local Ca^2+^ concentration in dendrites could be finely tuned, resulting in nearby synapses undergoing LTP or LTD, or left ‘protected’ from plasticity (Fig 5). These effects of synaptic inhibition on local dendritic excitability (modifying local dendritic NMDA and Ca^2+^ spikes) and, consequently, on synaptic plasticity are correlated with the impact of dendritic inhibition on burst activity. It seems that dendritic inhibition might simultaneously control local synaptic plasticity and global somatic burst activity (see also Owen, Berke, and Kreitzer 2018).

In the case of single-branch activation whole-tuft LTP is very robust to the location and timing of inhibition (Fig 5**d**1,3). Furthermore, different inhibitory locations generate varied plasticity maps: segmenting the branch to LTP and LTD (distal tuft inhibition; Fig 5**d**3), or to LTP, LTD and protection of weights (intermediate tuft inhibition; Fig 5**d**2). However, for the case of whole-tuft activation, all inhibition locations distal to the intermediate trunk with correct timing reduce the tree-wide LTP to highly localized LTD (trunk or distal inhibition; S5**c**1,3 Fig) or limited adjacent LTP (intermediate tuft inhibition; S5**c**2 Fig), making this burst class worse in producing extended LTP or diverse local plasticity.

We showed that pairing EPSPs in the tuft with somatic bursting generally produces LTP (Fig 5**d**1). This prediction is supported by several experimental results (Golding, Staff, and Spruston 2002; Letzkus, Kampa, and Stuart 2006; Kampa, Letzkus, and Stuart 2006) but disagrees with others (Birtoli 2004). In another paper, Owen et al. (2018) suggested a role for burst inhibition that is compatible with our findings: limiting excitatory synaptic plasticity for efficient and stimulus-specialized implementation of learning. That SOM inhibition of bursts leads to ineffective plasticity was shown in the hippocampus (Royer et al. 2012) but not directly and causally in the cortex.

### Related studies

In their theoretical study, Shai et al. (2015) showed that the intraburst firing frequency is best approximated by a composite sigmoidal function of the number of basal, and apical synapses, with apical number modulating the threshold of basal number required for bursting. This simplification is challenged by our findings, because it doesn’t generalize to the new class of bursts generated by single tuft branch activation (Fig 2). Specifically, their results do not show bursting for tuft only input, even at high numbers of synapses. Our findings require additional nonlinearities to account first for NMDA spikes and then for the non-monotonic transition in frequency between burst classes. The non-monotony of intraburst frequency is expressed in the corresponding heatmap (Fig 2**b** right). First more synapses mean higher frequency, but then it means shifting to coincidence-burst with lower frequency (compare to Shai’s Fig 4a left). This discrepancy arises from our clustering of the apical synapses compared to their whole-apical dendrite distribution, leading to stronger interactions (more spikes, S2 Fig) and a basal-independent Ca^2+^ spike (S1**c** Fig and top green region in Fig 2**b**). However, our findings support Shai et al.’s result of bursting with basal only input (Fig 2; see also Birtoli 2004), challenging the notion that bursts depend on Ca^2+^ spike firing (Helmchen et al. 1999; de la Peña and Geijo-Barrientos 1996).

Combined with our burst classes findings, we predict that input patterns generating coincidence many-spikes high frequency bursts (in which suppression of plasticity is more common) will also activate local SOM+ interneurons, and they would supply feedback inhibition near Ca^2+^ hotspot to withhold excessive LTP (Naud and Sprekeler 2018). This could be tested by recording SOM-L5PC pairs verified for feedback inhibitory connections, possibly with excitation of presynaptic axonal projections. Analyzing their activity during L5PC burst/BAC-firing and using long-term plasticity indicators (e.g., AMPA/NMDA ratios, spine sizes) would validate our predictions.

### Implications of burst control on perception

Larkum et al. (2009; 2013) suggested that the BAC firing mechanism implements coincidence detection between bottom-up sensory stream and top-down context modulation. Other studies made even stronger claims implicating bursts to be involved, via BAC firing, with conscious perception (Anastassiou and Shai 2016), visual illusions (Bachmann 2015) and attentional modulation of activity (Anderson, Mitchell, and Reynolds 2011). In view of our findings of two types of bursts, these effects could be reexamined, and the impact of dendritic inhibition of L5 pyramidal neurons on such high-level processing explored. Particularly, Takahashi et al. (2016; 2020) showed that optogenetic stimulation of L5bPC dendritic tufts enhances tactile stimuli detection, whereas blocking these cells’ outputs to subcortical target regions suppressed stimuli detection. We predict that distal tuft inhibition (e.g., by feedback via Martinotti cells) will attenuate local dendritic excitability and, in turn, decouple between the bottom-up and top-down input streams and, thus, disable conscious perception. As overreaching as the claims mentioned above seem to be, it is exciting to connect biophysical mechanisms such as specific dendritic inhibition to our subjective experience.

## Methods

All simulations were run using NEURON 7.7 (Hines and Carnevale 1997) and Python 2.7.16 (NumPy v1.16.2), initially on Windows/Linux PC, and for final plots on parallel processing cluster unit.

Code for obtaining the main results is available at https://github.com/EilamLeleo/burst. Additional data files are available on request from E.G.L.

### Model cell

We used the established compartmental model of thick-tufted L5bPC developed by Hay et al. (2011), including modifications of voltage gated calcium channel densities as in Shai et al. (2015) and I_h_ channel density distribution as in Labarrera et al. (2018). The two similarly functioning reconstructed morphologies were used to verify our findings (see Fig 6 in Hay et al. 2011), though plots were generated with the first for convenience and consistency.

Viewing our simulated voltage traces, we noted a biologically unrealistic amplitude and prolonged duration of after depolarization (ADP) in the somatic Na^+^ spikes. Scanning our parameter space, we decided to change a single maximal conductance value, so as not to significantly affect our results and the main findings of previous publications – that of the somatic calcium dependent potassium current (SKv3_1). The peak conductance value of the SK channel was thus increased by 1.5-fold compared to that of Hay et al. (2011) such that somatic g_SK_ = 3380*1.5 = 5070 pS/μm^2^.

### Simulations

Electric activity of the neuron was simulated for 600 ms at each instantiation with simulation dt = 25 μs. Simulation temperature was 34°C as previously suggested (Markram et al. 2015), and initial voltage was -76 mV.

We save all simulation data to Python NumPy arrays, initializing simulation of any experiment with each parameter combination at 100/200 random synaptic instantiations using a parallel processing unit (cluster). Randomly drawn properties were both spatial – site of synapse impinging on the dendrite, and temporal – activation time, as described in the following Input Distributions subsection. For most extensive data summarizing plots, we save only inter-spike intervals (ISIs), for calculation of spikes per burst and firing rate.

### Synapse models

Excitatory synapse model chosen (ProbAMPANMDA2.mod), implemented by Ramaswamy et al. (2011) and modified by Hay et al. (2011), combines a fixed ratio (of equal weights) of fast-AMPA (decay time constant τ_AMPA_ = 1.7 ms) and slower-NMDA (decay τ_NMDA_ = 43 ms) ionotropic receptors. Reversal potential for both was e = 0 mV. V_rest_ = -80 mV. Peak conductance gmax was fixed at 0.4 nS for both AMPA- and NMDA-synaptic conductances.

Inhibitory synapse model (ProbGABAAB_EMS.mod; Ramaswamy 2011) was preserved to include fast- decaying GABA_A_ only (decay τ_GABA_A_ = 8 ms), by keeping GABA_AB_ ratio = 0. Reversal e_GABA_A_ = -80 mV; with peak synaptic conductance g_max_ = 1 nS.

### Input distributions

Synaptic locations and activation times were randomly drawn from spatially uniform and temporally normal distributions, separated for basal and apical (excitatory) populations, and for inhibitory. For simplicity and lack of additional preliminary findings, all temporal distributions of any single simulation are identical in variance.

We note three main types of input patterns by how they spread on the dendritic tree: basal, apical or both. Basal branches are plentiful, interact at the soma and are less prone to NMDA-spike generation. Of course, some may be active spontaneously on the same branch and would contribute to an NMDA-spike formation, and a few of those will allow swift somatic APs or a burst (A. Polsky, Mel, and Schiller 2009). However, the apical tuft allows this more readily by having long thin branches that create NMDA-spikes by fewer synapses (Poleg-Polsky 2015), and by the Ca^2+^ hotspot transforming these into prolonged depolarizations at the soma. This lower threshold also means uniform distribution of synapses on the tuft will cause a burst before it would on the basal tree. Nevertheless, a bAP from the soma will lower the threshold for Ca^2+^-spike firing by the tuft and a burst, so the main options for bursting are tuft alone or both tuft and basal. How feasible, abundant and distinguishable are both input types? We show they are very much so.

For generating Fig 1 and Fig 2 and S1 Fig, excitatory synapses were scattered on the entire basal tree, and on a randomly drawn continuous 750 μm stretch of the apical dendritic tuft, which is about 1/10 of the entire apical length or 1/5 of the tuft - equivalent to a single offshoot of the tuft from nexus to all distal terminals. In Fig 1 and S1 Fig (50 basal and 30 tuft synapses) temporal distribution was varied in standard deviation σ from 0 (instantaneous) to 10 ms, with Δt ≠ 0 (delay between apical and basal mean activation times) introduced in Fig 1. For Fig 2 σ = 10 ms – comparable to Shai et al.’s (2015) 100 ms wide uniform distribution, while basal and apical synapse number were manipulated independently. The delay described by Larkum et al. (1999) from somatic spiking to apical EPSP isn’t the same as from basal to tuft synaptic activation, but we note a ∼3 ms delay of both – somatic spiking from mean basal activation and apical EPSP peak from tuft activation, so this equivalence is accounted for. Synaptic noise was introduced for in-vivo like state by input of the same magnitude, but uniformly distributed on the dendrites and in simulation-time (600 ms starting 100 ms before targeted synaptic activity). Results were insignificantly different (except for a minor reduction in apical synapse number threshold for bursting in Fig 2 from 50 to 40, not shown).

For plotting Fig 3-5, inhibitory synapses were introduced on the apical tree. In all three, they were manipulated by distance from soma (20 synapses in < 200 μm disparity). Temporal jitter was allowed with the same parameters as excitation, excitation on both trees were activated simultaneously (Δt = 0), and Δt_inh_ was introduced to separate between excitation to inhibition mean activation times. For Fig 3 and Fig 5, the number of excitatory synapses was increased from 50 basal and 30 tuft in Fig 1 to 70 basal and 40 tuft synapses, owing to the higher bursting threshold at σ = 10 ms. Tuft excitation was restricted to a single continuous branch from nexus to the most distal terminal, inhibition is only on the trunk or the same branch. In Fig 4 excitation is on the entire tuft, and inhibition is at a fixed distance, but in all oblique or tuft branches sharing this distance from the soma.

### Data analysis

Most initial and direct analysis of electrodynamics in the model L5PC was calculated online on Python after each run of NEURON. Gathering of all data in any single experiment for plotting results and drawing conclusions was generally executed manually.

### Spike detection

We set the spike detection somatic voltage threshold at 0, though it essentially does not differ by setting it at -20 mV or +20 mV, nor by detecting it from the voltage trace at the axon initial segment. Spike width did not exceed 2 ms, and minimal ISI was above 3 ms, so effectively the soma was instantly hyperpolarized below detection threshold right after spiking, thus allowing for the next to be detected even at high intraburst (between spikes in a burst) firing rates of ∼300 Hz.

### Number of spikes and firing rate calculation

The definition of bursts relates to statistical deviation of firing rate from random Poisson firing. A cell is considered bursting if either the coefficient of variation (CV = standard deviation/mean) or the Fano factor (= variance/mean) of ISIs over some time interval, is higher than 1 (Poisson). Relying on mean firing rates of pyramidal cells, we allow ourselves to group spikes as part of a burst by their ISI alone, < 20 ms (> 50 Hz): A burst of > 50 Hz will occur by chance for a neuron with a high 5 Hz mean firing rate (FR) approximately once every 10^15^ seconds. So, we set ISI ≤ 20 ms as our burst-grouping interval, and we note that our generated cell intraburst FR exceeds 100 Hz significantly. If no two spikes arrive within 20 ms of one another, then we count only one spike.

By this criterion we count and average the number of spikes per burst (at each simulation instance), even if isolated spikes precede the burst. The Δt range in Fig 1**f** is defined by > 95% chance for bursting and shown in Fig 1**d** as > 2 spike per trial average. Firing rate in Fig 2**b** and 2**c** is the mean over all spikes of all bursts in the same parameter combination, taking isolated spikes as a 0 Hz burst. Measuring intraburst FR for < 2 mean spikes per burst seems peculiar, because at least 2 spikes are needed in order to measure it, still it proves useful to keep values not only from bursts. We tested for big variations inside a single burst, that would discredit our conclusions – a burst of one ISI at 4 ms and another at 6 ms is not firing at 200 Hz but rather in the 170-250 Hz range. We find no substantial (1.5- fold) differences occurring in over 5% of instances at any parameter combination.

### Generating plots

All voltage traces represent a single representative simulation. Input pattern was kept fixed for parameters not probed at the particular experiment, i.e., changes in activation times do not alter spatial locations of synapses in Fig 1 and S1 Fig, and variations in inhibition do not modify excitatory pattern in Fig 3-5. In contrast, each value on the summarizing graphs and heatmaps (in Fig 2) averages 100-1000 random spatiotemporal synaptic patterns, with matching distribution parameters. Error bars are missing from all graphs that plot more than a single line. In Fig 1**f** and 1**g** they reflect sampling resolution. In calculating mean spikes per burst, variance values are large, as any mean close to half a whole spike would indicate > 0.25 variance. This will not reflect the consistency of our measurement, so instead we use (in Fig 3**d**) the variance of batch means (10, each from 20 repetitions). Hence, the values are very generalizable and will occlude a possible variation within the fixed-parameter data.

The scatterplot in Fig 2 transforms the input-aligned picture on the heatmaps, into a mapping of each mean spike per burst value with its corresponding FR (each point represents a spike # from Fig 2**b** left and its collocated FR to the right). There we directly assess their correlations, and cluster output parameters pairs to three dense regions, high-rate low-spike #, low-rate high-spike # and zero spikes. These clusters in-turn map to correlating histograms, showing both parameter ranges attributed to each kind, when separated by input synapse numbers: orange generated by > 40 tuft and < 120 basal synapses; blue by > 150 basal and < 100 tuft synapses.

### Calcium dependent plasticity

The learning rule applied is summarized by the two equations below (Eq. (1) and (2)) and graphs (Fig 5**a**) shown. The equations define the plasticity function Ω and the resulting learning rule as the change in synaptic strength (w_j_). θ_p,d_ are concentration thresholds for potentiation and depression respectively (Fig 5**a**1 and 5**c**), and η the learning rate (Fig 5**a**2).

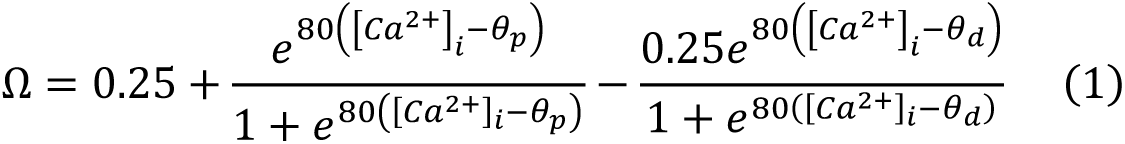

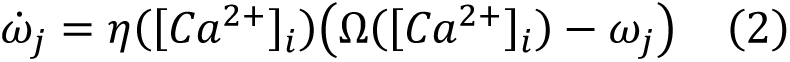

Graphs are created for maximal [Ca^2+^]_i_ (concentrations) during the simulation at different inhibitory locations and times (Fig 5**c**). Threshold was slightly shifted from the previous studies (Shouval, Bear, and Cooper 2002; Bar-Ilan, Gidon, and Segev 2013) to θ_p_ = 0.4 μM, θ_d_ = 0.25 μM (originally 0.5 and 0.3), which creates a more variable and realistic range of phenomena (LTP/D & PROT.). To generate synaptic weight changes in Fig 5**d**, the duration over or under any threshold is multiplied by a learning factor chosen as ten repetitions for effect size, as physiological [Ca^2+^]_i_ measurements are difficult to control for in these relatively short simulations. Synapses on trees represent predicted weights after learning by this rule.

## Acknowledgements

We would like to thank David Beniaguev, Michael Doron, Oren Amsalem, Hadas Manor, Yair Deitcher, Tal Nahari and all members of the Segev lab for many insightful discussions and valuable feedback. This work was supported by the generous support of the Drahi family foundation, a grant from the European Union’s Horizon 2020 Framework Program for Research and Innovation under the Specific Grant Agreement No. 785907 (Human Brain Project SGA2), the ETH domain for the Blue Brain Project, the Gatsby Charitable Foundation, the NIH Grant Agreement U01MH114812.

## Supporting Information

### S1 Appendix. Additional burst control

Input to the apical obliques was suggested to better couple input to the apical tuft with that to the perisomatic/basal compartments, which is beneficial for BAC firing (M. E. Larkum 2004). Our simulations show no significant role for these synapses.

A different dendritic spiking mechanism than BAC firing is driven by the coactivation of clustered NMDA receptors in neighbor dendritic synapses. Having a longer time scale of activation (large time constant) than AMPA receptors, higher conductance values, and high penetration to Ca^2+^ (A. Polsky, Mel, and Schiller 2009), NMDA synapses may largely dictate the neuronal response. But in large pyramidal cells, some of these synapses are located on the far distal tuft, and hardly give rise to a somatic excitatory post-synaptic potential (EPSP) due to voltage attenuation along the elongated apical dendrites (A. Polsky, Mel, and Schiller 2009). Therefore, what is their function?

A considerable number of activated tuft synapses, especially in conjunction with a bAP, may ignite a Ca^2+^ spike and a burst. Besides, NMDA synapses show voltage dependence, which enhances the coactivation of others nearby by relieving channel blocking Mg^2+^ ions. This dependence creates the NMDA spike, which requires less coactivated synapses in the tuft than in basal dendrites (Poleg-Polsky 2015). This phenomenon appears either isolated in a dendritic fragment of ∼30-70 μm (focal) or in a global whole-tuft reach. Actually, researchers have claimed that NMDA spikes in the tuft directly and significantly affect somatic firing only by generating a Ca^2+^ spike (A. Polsky, Mel, and Schiller 2009; Grienberger, Chen, and Konnerth 2014). We showed that activating some tens of AMPA/NMDA synapses on a moderate length of dendritic tuft (350-750 μm) creates extended NMDA spikes, initiating a Ca^2+^ spike (or decoupled from it by inhibition) and a burst of fewer spikes and higher rate than that of BAC-firing (Fig 2).

To better simulate natural input by decreasing temporal input correlations, we introduced a normal distribution for drawing activation times, and gradually increased its variance (σ^2^; Fig 1**b** and S1**b** Fig). For examination of tuft-to-basal mean activation delay we separated the distribution in two of identical σ^2^ and different means, denoting their disparity Δt. This distribution form allows dependence on a variable separation between two distributions, as opposed to an overlap range only in uniform distributions. A supralinear sum of simultaneous synaptic activations would decrease if they were distributed over longer timescales (S1**c** Fig) due to reduced overlap. This dispersing is equivalent to decreasing the number of active synapses at a fixed time interval. Consequently, the dendritic event amplitude and spikes per burst, or bursting altogether, may diminish (Fig 1**c** and S1**c** Fig). Raising σ values from 0 (instantaneous) to 3 ms did not change the output burst significantly.

Typical *in vivo* conditions may create stronger spatial correlations (i.e., clusters of neighbor synapses) and weaker temporal correlations (dispersed activation times). So, we continued by checking the optimal spatial extent of synapses in the tuft for bursting (S2 Fig). Focusing first on tuft extent is partially justified because the basal dendrite requires more synaptic activations for NMDA spikes (Poleg-Polsky 2015), affords less proximity to Ca^2+^ hotspot, and interactions between branches are obtained at the soma and not on dendrites. We found the highest burst probability and spikes per burst arise for synapses distributed over 350-750 μm of a continuous dendritic length (S2 Fig, as illustrated in Fig 1**a**) which is optimal for supralinear summation and does not begin to saturate (by a few tens of synaptic activations) the membrane voltage. Synaptic activation confined to a smaller dendritic extent or spread over larger areas will decrease spike number.

**Fig. S1:**
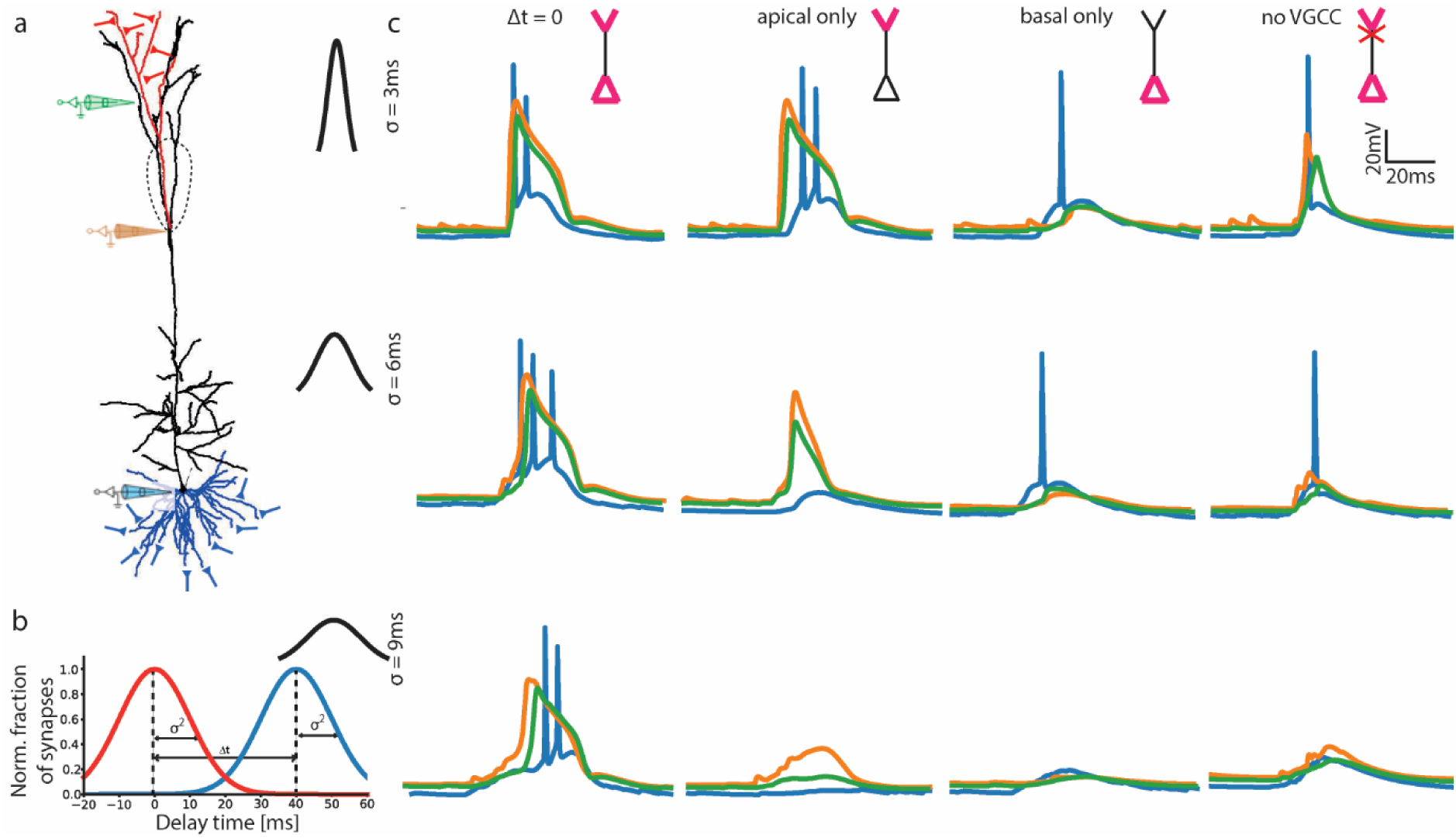
Burst initiation by basal and tuft input distributed in time. **a**. Model L5PC, schematically showing 50 basal (blue) and 30 apical tuft (red) excitatory synaptic locations (drawn from uniform distribution over the appropriate colored region). Electrodes are shown for corresponding voltage traces in **c**. **b**. Temporal distribution of input. Activation time is drawn from two tree-specific (same colors as **a**) normal distributions with equal variance and mean (unless noted otherwise). **c**. Voltage traces showing outcome of input to basal/tuft/both trees, with three σ values and no-VGCC control. Coincidental input to both trees results in burst firing for all σs. Tuft activation alone creates a burst for small σ, a dendritic spike for intermediate and nothing for high (equivalent to a lower synapse number). Basal activation results in a single AP at most, and removing VGCC returns all to one spike.

**Fig. S2:**
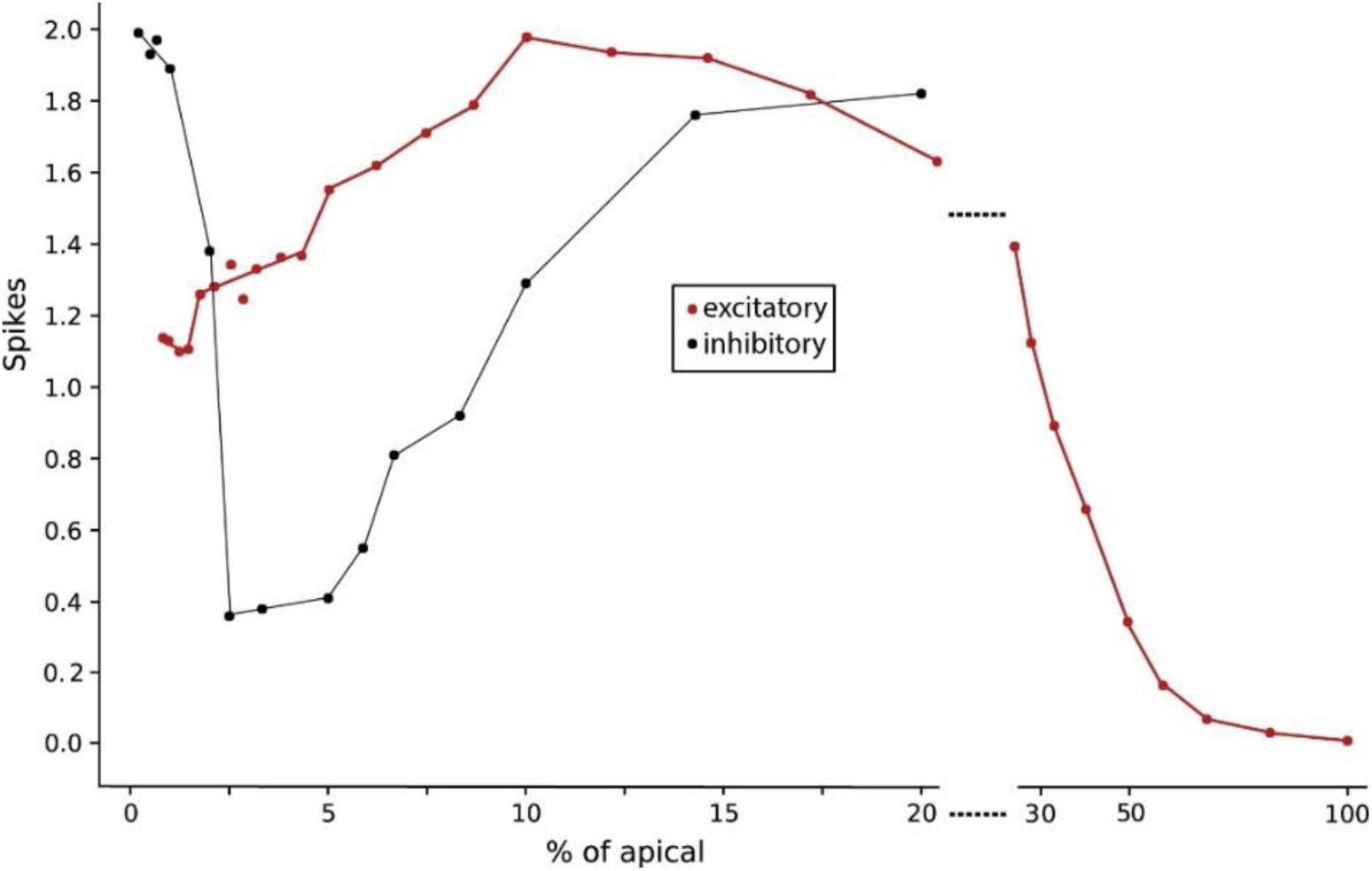
Spatial extent of synapses controls burst generation. Spikes generated as a function of percent of apical tree on which excitatory alone (red) or inhibitory (black; excitation on 10%) synapses are distributed. Excitation continues over 20% at a different scale. Excitatory conditions as in Fig 1 (σ = 9 ms, Δt = 0), inhibition as in Fig 3.

**Fig. S3:**
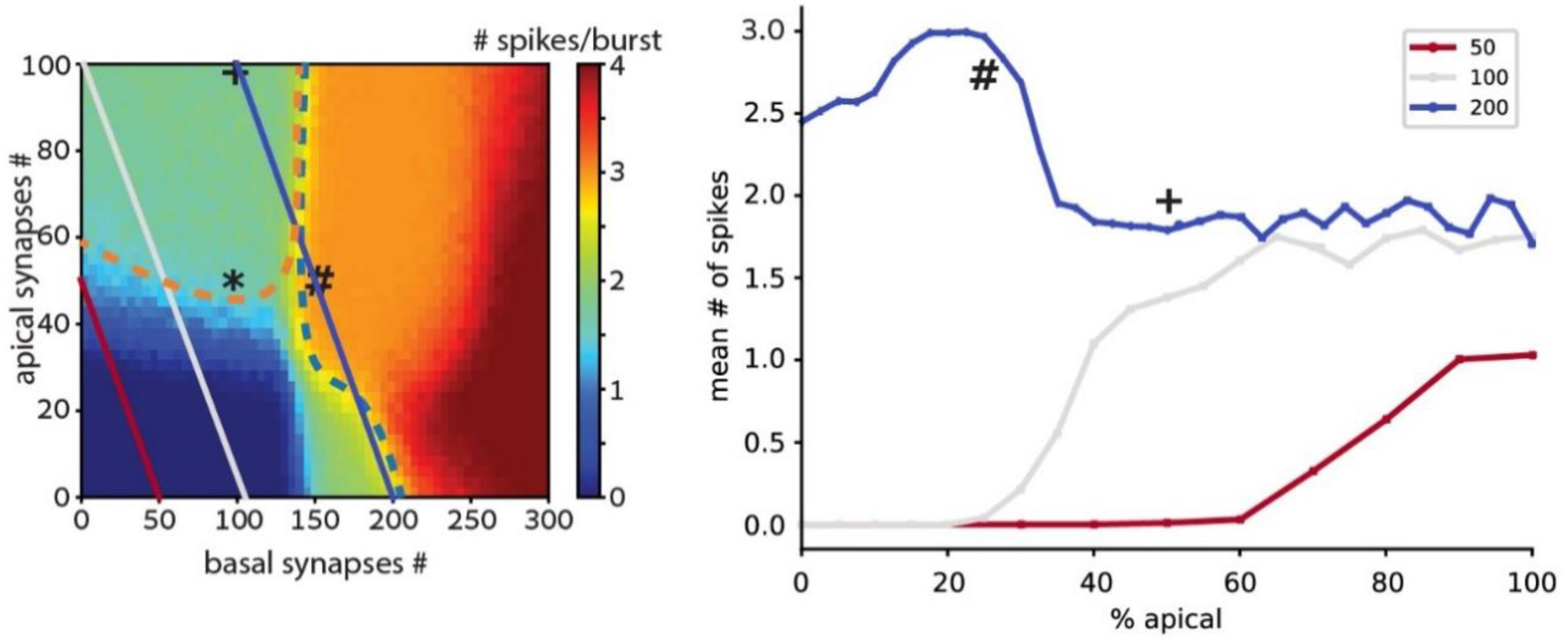
The number of spikes per burst with fixed total activations depends on apical:basal ratio. Left: heatmap of mean number of spikes per burst as a function of the number of activated synapses on the basal and apical trees, from Fig 2. Overlaid are fixed total synapse number diagonal lines, whose profile is plotted to the right. Right: Mean number of spikes per burst, as a function of the ratio between apical and basal synapse number, for various total synapses, as plotted on the respective colored diagonals on the left. σ = 10 ms, Δt = 0.

### S2 Appendix. “Off-path” inhibitory control of bursting

Inhibition at or close and distal to the Ca^2+^ hotspot (“off path”, similarly to Gidon and Segev 2012) disrupted the Ca^2+^ spike and burst from forming (Fig 3**b** and 3**c**). These results relate to previous works showing that distal inhibition (“off path”) is more efficient at terminating dendritic spikes than proximal inhibition (“on path”; Gidon and Segev 2012). They keep all excitatory input in a single point (“hotspot”) able of generating a Ca^2+^ spike, whereas we distributed the excitation, meaning that our “off path” is actually between the majority of excitatory synapses and the Ca^2+^ hotspot, rather than distal to them. Distal tuft inhibition still better attenuates nearby NMDA spikes but not the Ca^2+^ and Na^+^ spikes directly. Somatostatin-expressing (SOM) interneurons were found to innervate L5PC tuft (Y. Wang et al. 2004), and within it the Ca^2+^ hotspot. Thus, we draw the conclusion that regulation of burst firing is amongst their core roles (similarly to Royer et al. 2012; Goldberg et al. 2004).

### S3 Appendix. Complementary plasticity comparisons

**Fig. S4:**
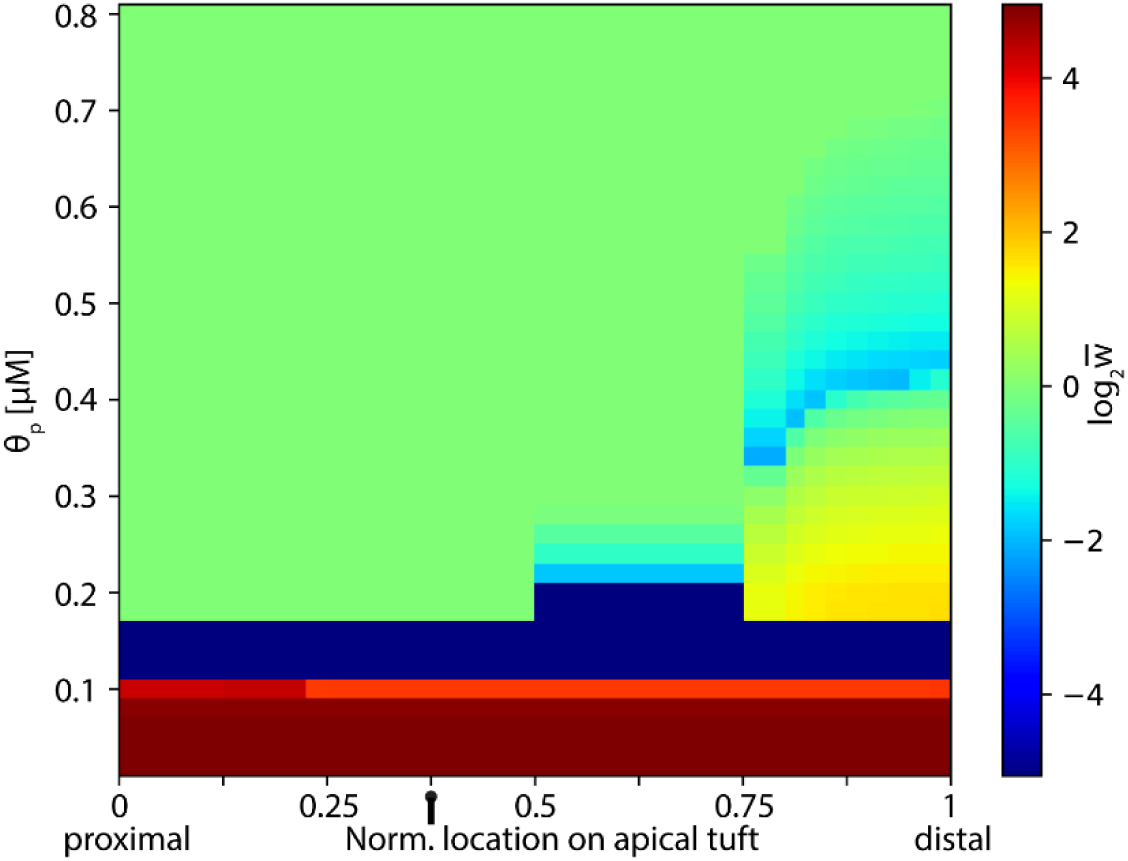
Ca^2+^ dependent plasticity patterns at different thresholds. [Ca^2+^]_i_ measured as in Fig 5c2 (intermediate inhibition location, Δt = 0) were used for calculating plasticity modifications as in Fig 5c2, for a range of threshold θ_p/d_ values. The heatmap represents synaptic weight after plastic modification, normalized to initial value and colored on a log_2_ scale (Red LTP, blue LTD, 0 – green protected). Modified excitatory synapse location is ordered on x-axis. LTP threshold θ_p_ on y-axis. LTD threshold θ_d_ is kept at a fixed ratio of 0.6 to θ_p_.

**Fig. S5:**
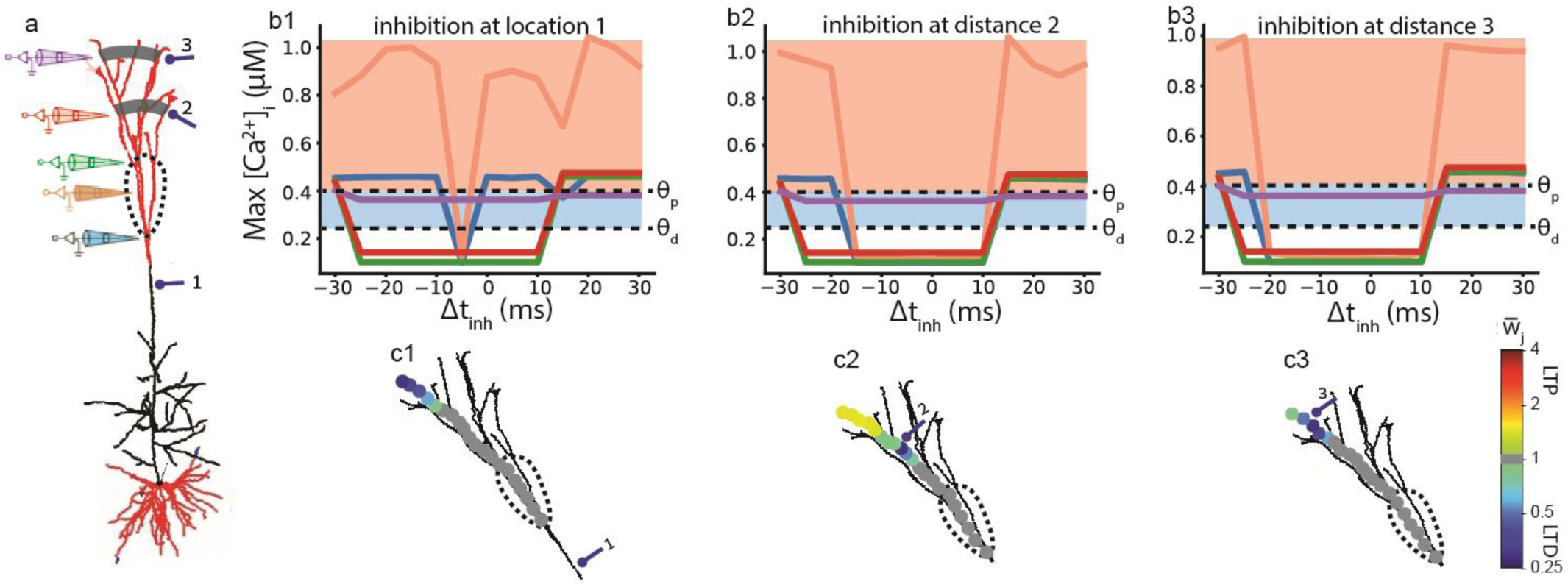
Ca^2+^-based plasticity of excitatory synapses during burst suppression in the whole-tuft case. **a.** Model neuron as in Fig 4a. Red branches are excited. Electrodes for [Ca^2+^]_i_ measurement as in Fig 5a. Inhibitory synapses at all locations of a fixed distance from the soma (a single numbered grey shaded strip). Dashed line denotes Ca^2+^ hotspot. **b1-3.** Maximal [Ca^2+^]_i_ along the apical dendrite at the three locations (colored electrodes in b) as a function of Δt_inh_ between excitation and inhibition for each inhibition location. Dashed lines mark thresholds, shadings the resulting change - LTP (red), LTD (blue). **c1-3.** Synaptic weights after repeated execution of the learning rule (Eq. (1), Methods) for 10 input repetitions (60 seconds) and Δt_inh_ = -5 ms at each inhibition distance # **(a,b)**. Twenty representative excitatory tuft synapses are color plotted by their respective weights: yellow - LTP (in 2), blue - LTD, and gray - protected.

Our previous results imply diverging outcomes for plasticity by the different burst classes and corresponding inhibition. We test this by applying the same Ca^2+^-dependent plasticity procedure, while using the whole-tuft excitation configuration as in Fig 4. The results are well in line with our hypothesis, as full tuft excitation is mainly associated with lower [Ca^2+^]_i_ than the single-branch case, which generally means reduced plasticity effects in the burst-suppressed dendritic tuft (S5 Fig). In response to bursting, without inhibition, our model produces strong LTP at the hotspot, and minute LTP throughout the apical tuft (on average, although some sites may show LTD). We defined inhibition distances at 500, 1100, and 1300 μm from the soma for comparison with the single-branch case (Fig 5). We remind that in this synaptic configuration inhibition is distributed in all branches at these distances, and note the difference from same distance indexes in Fig 4. Distal tuft inhibition (#2-#3) activated even 20 ms before or synced to excitation (as in Fig 4) drops [Ca2+]i below the effective thresholds (S5b2, b3 Fig), resulting in protected synaptic efficacies, with the exception of the most distal branches which are depressed (S5c3 Fig) or slightly potentiated (S5c2 Fig).

Trunk inhibition (location #1 in S5a Fig) too shows suppression of the Ca^2+^ spike and burst, and minor distal LTD, but adheres to a highly specific timing restriction, limited to inhibition 5 ms prior to excitation (dip in orange and blue lines in S5b1 Fig, and see Fig 1; Matthew E. Larkum, Zhu, and Sakmann 1999). This timing supports both the bAP role in burst initiation (identifying our coincidence class), and vicinity to the bursting threshold, because no other delay suppressed bursting.

Such complex multilevel simulations as our plasticity experiments, presume many assumptions, some of which we did not tackle explicitly (dominance of [Ca^2+^]_i_ in plasticity, learning rule shape thresholds and saturation, dendritic shaft equivalence to spines, influx through VGCC and not NMDAR, no diffusion, etc.). Even so, it allows an elaborate demonstration, starting from the established effect of global dendritic Ca^2+^ spike that promotes LTP throughout the tuft (Fig 5e1; Golding, Staff, and Spruston 2002). Building on this foundation, precise inhibition will suppress either the local NMDA spike or the widespread Ca^2+^ spike, thus lowering [Ca^2+^]_i_ at surrounding sites and effectively weakening or protecting synaptic weights (Fig 5e1-3).

A parallel line of research suggests Ca^2+^ spiking as a means of information multiplexing. This multiplexing requires decoupling of tuft and perisomatic input, the tuft determining burst-probability and the perisomatic the event (burst or spike) rate (Naud and Sprekeler 2018; Payeur, A., Guerguiev, J., Zenke, F., Richards, B., & Naud 2020). Balduzzi & Tononi (2013) argue that an efficient information coding scheme must emphasize selective responses with bursts. An elementary experiment we ran involved training a convolutional neural network on reproducing I/O pairs of our detailed model, and revealed bursts as highly specialized responses, explained by significantly fewer principal components (Beniaguev, unpublished data).

